# NanoPlasmiQC: Full plasmid sequencing with ONT long-reads and automatic data analysis

**DOI:** 10.64898/2026.04.01.715842

**Authors:** Julie Anne V. S. de Oliveira, Vinson Ng, Katharina Wolff, Boas Pucker

**Author notes:** contributed equally (Julie Anne V. S. de Oliveira, Vinson Ng).

## Abstract

Long-read sequencing has shown rapid technological development in recent years. It has been established as the standard method for the sequencing of plant genomes and has also gained importance for full plasmid sequencing. As Sanger sequencing has a limited read length of about 1 kb, long-read sequencing offers a great advantage, as the full plasmid can be sequenced in one read.

Here, we present a cost-effective workflow to sequence full plasmids and compare the results against an expectation. The per-plasmid cost of this workflow is determined by the number of plasmids investigated simultaneously, but can be lower than the price of a single Sanger sequencing reaction. We developed a workflow for automatic data processing, which allows us to complete sequencing and data analysis within a day.

## Introduction

Since the invention of chain termination sequencing (Sanger *et al*., 1977), often referred to as ‘Sanger sequencing’ to honor the inventor, researchers have benefited greatly from unraveling the sequence of DNA molecules. For almost 50 years, sequencing technologies have shown rapid development, with massive parallel sequencing of the second generation and long reads of the third generation representing the most important breakthroughs (Margulies *et al*., 2005; Metzker, 2010; Mardis, 2013, 2017; Pucker *et al*., 2022; de Oliveira *et al*., 2026). In plant genomics, the long-read technologies offered by Pacific Biosciences (PacBio) and Oxford Nanopore Technologies (ONT) are the most important sequencing methods (Marks *et al*., 2021; Schwacke *et al*., 2025; de Oliveira *et al*., 2026). In addition to numerous applications in agriculture (Zhou *et al*., 2020; Jayakodi *et al*., 2020; Walkowiak *et al*., 2020; Pucker *et al*., 2022), plant genome sequences are fundamental to unraveling complex biosynthesis pathways leading to products with biotechnological or biomedical value (Cheng *et al*., 2025; Hakim *et al*., 2025).

Nanopore has also frequently been used to characterize the products of mutagenesis or genome editing experiments to identify any off-target effects. T-DNA insertion lines, generated by randomly inserting a T-DNA into a plant genome, were fundamental for plant biology research because homologous recombination does not work effectively in plants. While these lines form an important resource for the discovery of gene functions, a comprehensive genetic characterization is necessary to rule out additional events besides the desired gene knock-out. An efficient way to characterize such lines is long-read sequencing combined with a local assembly for all T-DNA insertion loci (Pucker *et al*., 2021). A recently released pipeline allows the automatic analysis of CRISPR/Cas editing sites based on long reads (Chen *et al*., 2026).

Assessing gene functions and biotechnological studies often involve the construction of plasmids. For many years, plasmids were validated by Sanger sequencing. It is believed that up to 40% of certain plasmid types with inverted tandem repeats might contain an unexpected mutation (Bai *et al*., 2025). Circuit-seq was developed to analyze plasmids with nanopore sequencing without prior knowledge about the plasmid sequence (Emiliani *et al*., 2022). Until recently, plasmid sequencing relying only on long reads was not recommended due to small insertions/deletions caused by noisy long reads (Hernandez *et al*., 2024). However, with recent technological improvements leading to a raw read accuracy of about 99 %, full plasmid sequencing utilizing nanopore long reads could be feasible now. Especially ambitious projects involving large plasmids adopted full plasmid sequencing as an attractive alternative to combining numerous Sanger sequencing reactions. Full plasmid sequencing removes the need to design sequencing primers and ensures that the complete plasmid, and not just the newly inserted DNA construct, is checked. Costs of commercial offers for full plasmid sequencing often exceed 10 € per plasmid. Given that many biotechnological projects involve the generation and validation of numerous plasmids, costs for the sequencing add up. Sophisticated pooling systems have been proposed to reduce these costs by mixing multiple plasmids into one sample without barcoding (Uematsu & Baskin, 2025).

Here, we present a workflow for cost-effective and sustainable full plasmid analysis based on ONT long-read sequencing. The steps of sample preparation, library construction, and nanopore sequencing are described. An automatic data analysis workflow implemented in Python generates user-friendly outputs that enable plasmid validation by life scientists.

## Materials and Methods

### DNA preparations

All plasmids submitted for sequencing should have a concentration of about 50 ng/µL (NanoDrop measurement) and should be resuspended in TE buffer. Although differences in plasmid size are going to cause different sequencing depths, this approach makes it convenient for users, and the total coverage should be sufficiently high to resolve even plasmids with a lower concentration. Plasmids are pooled together, taking 1µL of each plasmid. While an optimization would be possible based on the size of the different plasmids, this approach ensures efficient handling of large numbers of samples. To ensure that no problematic chemicals can interfere with the nanopore sequencing, the plasmid mix is subjected to DNA precipitation. About 50 µL of 1 M NaCl is added to the plasmid mix. The precipitation is achieved by adding an equal volume of isopropanol to the mix of plasmids with NaCl. After centrifugation at >13,000 x g for 15 minutes, the supernatant is discarded. The sediment is washed with 70% ethanol. The precipitated plasmid DNA is then resuspended in 20 µL of water. The plasmid concentration is measured and diluted to a concentration of 10-15 ng/µL, as a total of 150 ng is to be used in the library preparation.

### Long-read sequencing

The library preparation is done with 10 µL of pooled plasmids using the Rapid Sequencing Kit from ONT (SQK-RAD114). Sequencing is conducted on R10 flow cells on a PromethION after completion of plant genome sequencing projects. The nanopores recovered through a DNase-based washing step (EXP-WSH004) still offer sufficient capacity for the sequencing of dozens of plasmids. Sequencing is typically performed overnight to ensure sufficient data are available for the following analysis. In cases of insufficient sequencing output due to a significantly degraded flow cell, it is possible to recover the library and to transfer it to a new flow cell. Basecalling for the first runs was done with Dorado v1.4.0 (ONT) in high accuracy mode (HAC). Later plasmid sequencing runs utilized Dorado v2.0.1 and v2.1.0 in HAC or super high accuracy (SUP) mode.

### Data analysis

Expected sequences of all plasmids were collected in a FASTA file. Headers in this file were cleaned of invalid characters as part of the data analysis process. A read mapping with minimap2 v2.26-r1175 (Li, 2018) against all expected sequences was conducted with the flag secondary=no in addition to default parameters. The resulting mapping was split per plasmid reference using samtools v1.19.2 (Li *et al*., 2009), and the reads per plasmid were extracted and converted into a FASTQ file with samtools. Variant calling per plasmid reference was performed with bcftools v1.19 (Danecek *et al*., 2021) using default parameters. Filtering of the extracted reads was done with seqkit v2.3.0 (Shen *et al*., 2024) to ensure that no redundant reads were contained. Due to very high coverage, subsampling was required to reduce the number of reads subjected to the following assembly step. Miniasm v0.3-r179 (Li, 2016) was used for a de novo assembly per plasmid using default parameters. Racon v1.5.0 (Vaser *et al*., 2017) was applied to polish the assembled plasmid sequence. Integrative Genomics Viewer v2.7.2 (Robinson *et al*., 2023) was used for manual inspection of results. The entire data analysis workflow was wrapped into a Python script, which is available via GitHub (https://github.com/bpucker/NanoPlasmiQC).

## Results & Discussion

We designed a workflow to enable cost-effective and sustainable plasmid sequencing with ONT long-reads (**Fig. 1**).

**Fig. 1:**
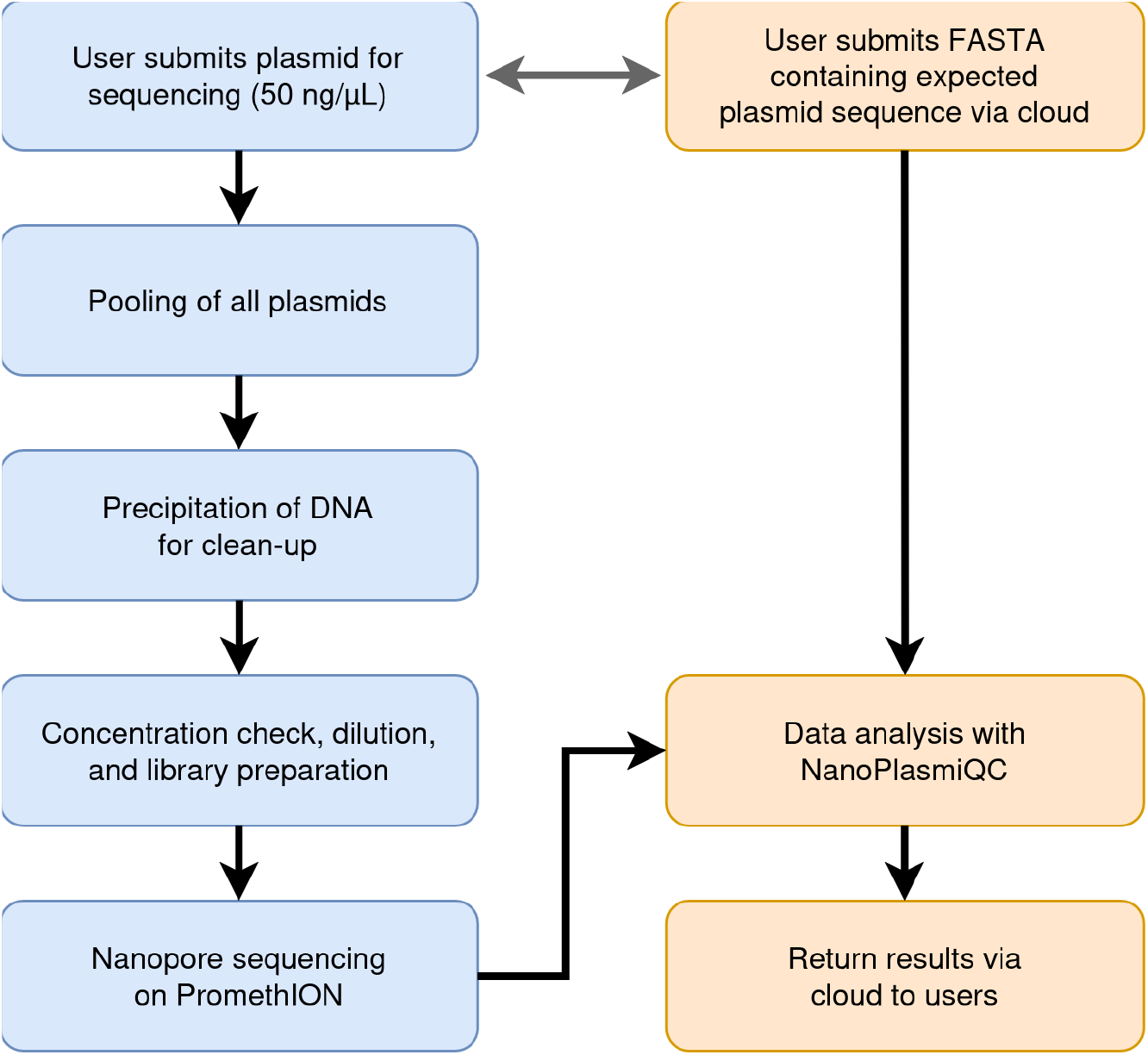
Workflow designed for full plasmid sequencing with ONT long-reads.

### Sequencing output

Sequencing a mixture of plasmids on a reused PromethION flow cell is expected to return low output when compared against typical plant sequencing runs. However, we managed to get sufficient sequencing data in our test runs (**Table 1**). Plasmids typically have a size of 5-20 kbp and can therefore be fully covered by individual reads. Usually, the limited size of plasmids results in a low N50 of <10 kbp. An average sequencing depth of 10x per plasmid should be sufficient to validate the sequence of most plasmids. Since the pooling of plasmids with different concentrations and sizes will result in biases, we aim for a coverage of >100x to ensure that even large plasmids are well captured in the resulting dataset.

**Table 1:** Plasmid sequencing output statistics. ^a,b^ The datasets generated in these runs have been merged prior to analysis to demonstrate that recovering a library from a flow cell for an additional sequencing run is possible.

| RunID | Output [Gbp] | N50 [bp] | Number of reads | Average read length [bp] |
| --- | --- | --- | --- | --- |
| RP080 | 12.75 | 7219 | 3405632 | 3745 |
| RP094 | 0.58 | 2584 | 487472 | 1199 |
| RP114<sup>a</sup> | 0.32 | 9991 | 163313 | 1928 |
| RP116<sup>a</sup> | 2.1 | 7666 | 391904 | 5335 |
| RP136<sup>b</sup> | 4.1 | 16.6 | 538447 | 7542 |
| RP137<sup>b</sup> | 1.7 | 16.7 | 219422 | 7956 |
| RP155 | 18.4 | 10629 | 2844374 | 6471 |
| RP171 | 9.2 | 10093 | 1316753 | 7009 |
| RP189 | 1.9 | 6913 | 333291 | 5995 |
| RP196 | 3.6 | 13858 | 649485 | 5581 |
| RP205 | 3.8 | 8989 | 766670 | 4968 |
| RP207 | 6.2 | 12471 | 8569217 | 718 |

Given the substantial sequencing output that can be achieved in most runs, plasmid sequencing offers the opportunity to include also slightly larger samples like microbial genomes. Per run, one bacterial, one archaeal, and one fungal sample can be included, because a de novo assembly and following separation of the sufficiently divergent species is possible.

### Implementation of the plasmid validation tool

Processing of the sequencing reads in order to validate the accuracy of constructed plasmids was automated with a Python script serving as a wrapper for numerous analysis steps (**Fig. 2**). The provided reference FASTA file is cleaned to ensure no problematic characters in the headers can break downstream tools (1). Statistics of the provided FASTQ file are calculated for documentation purposes (2). Minimap2 is run to align ONT long reads to the expected plasmid sequences with the secondary=no flag to ensure specific read mapping (3). Resulting SAM files are converted into BAM, sorted, and indexed (4). The BAM file is split by plasmid reference sequence with samtools to enable independent analysis of all plasmids in the downstream steps (5). For each BAM file, the reads are extracted in FASTQ format with samtools using the -F2308 flag and checked for duplicates with seqkit (6). The coverage per plasmid is calculated to enable subsampling to reach a target coverage (7). Based on the subsampled reads per plasmid, minimap2 and miniasm are used to de novo assemble the plasmid sequence (8). These assemblies are polished with Racon (9). The provided FASTA file with all plasmid reference sequences is also split accordingly and indexed with samtools (10). Variant calling is conducted with bcftools using the “-Q 10” and “-q 10” flags, as well as downstream filtering with QUAL>20 and DP> a coverage cutoff (11).

**Fig. 2:**
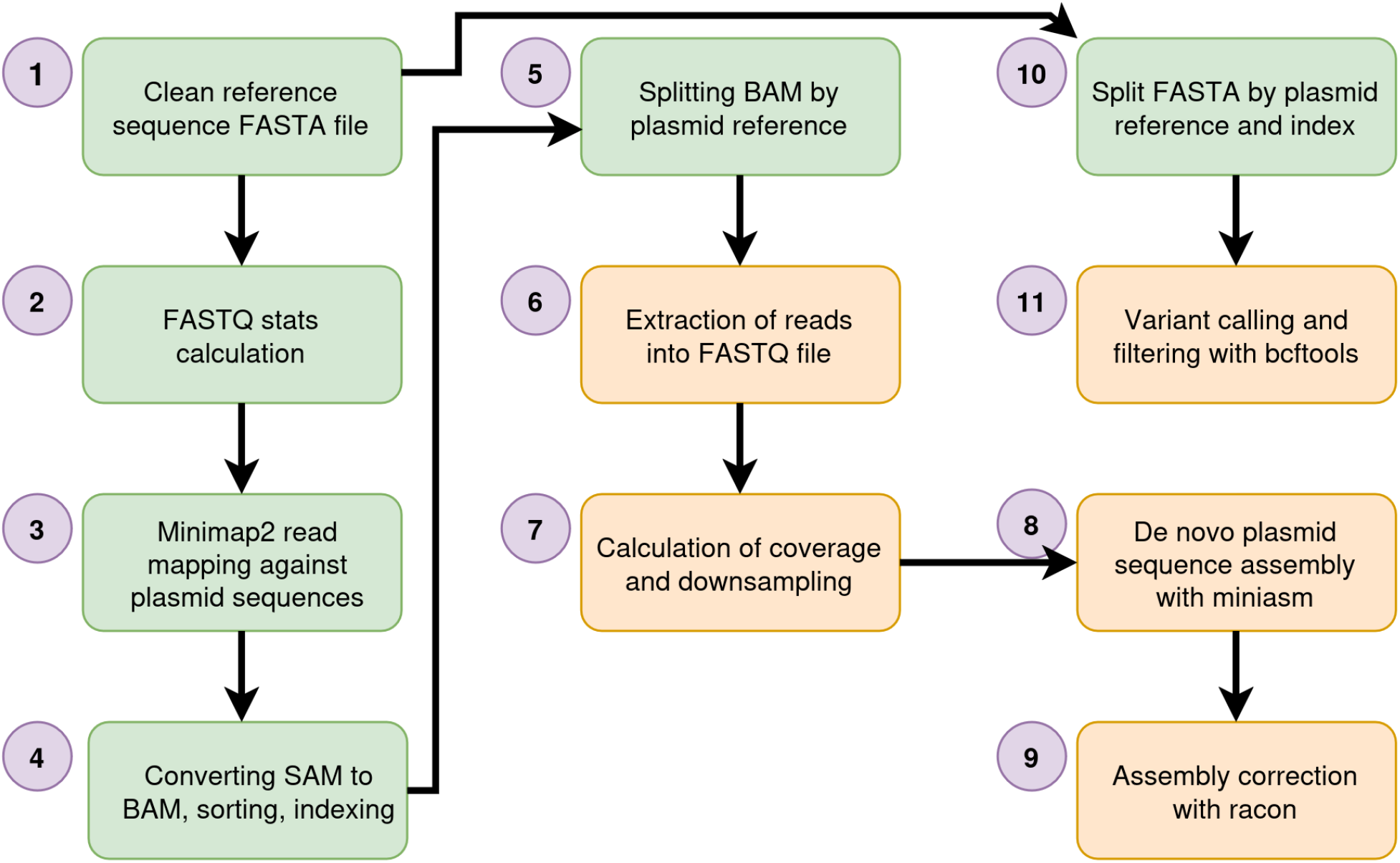
Plasmid sequencing data analysis workflow.

### Proof of concept

As a proof of concept, the previously published plasmid pBF3038 (Fang *et al*., 2011) was sequenced and assessed with the presented workflow. To illustrate the results obtained through the automatic data analysis, the results of pBF3038 were visualized with IGV(**Fig. 3**).

**Fig. 3:**
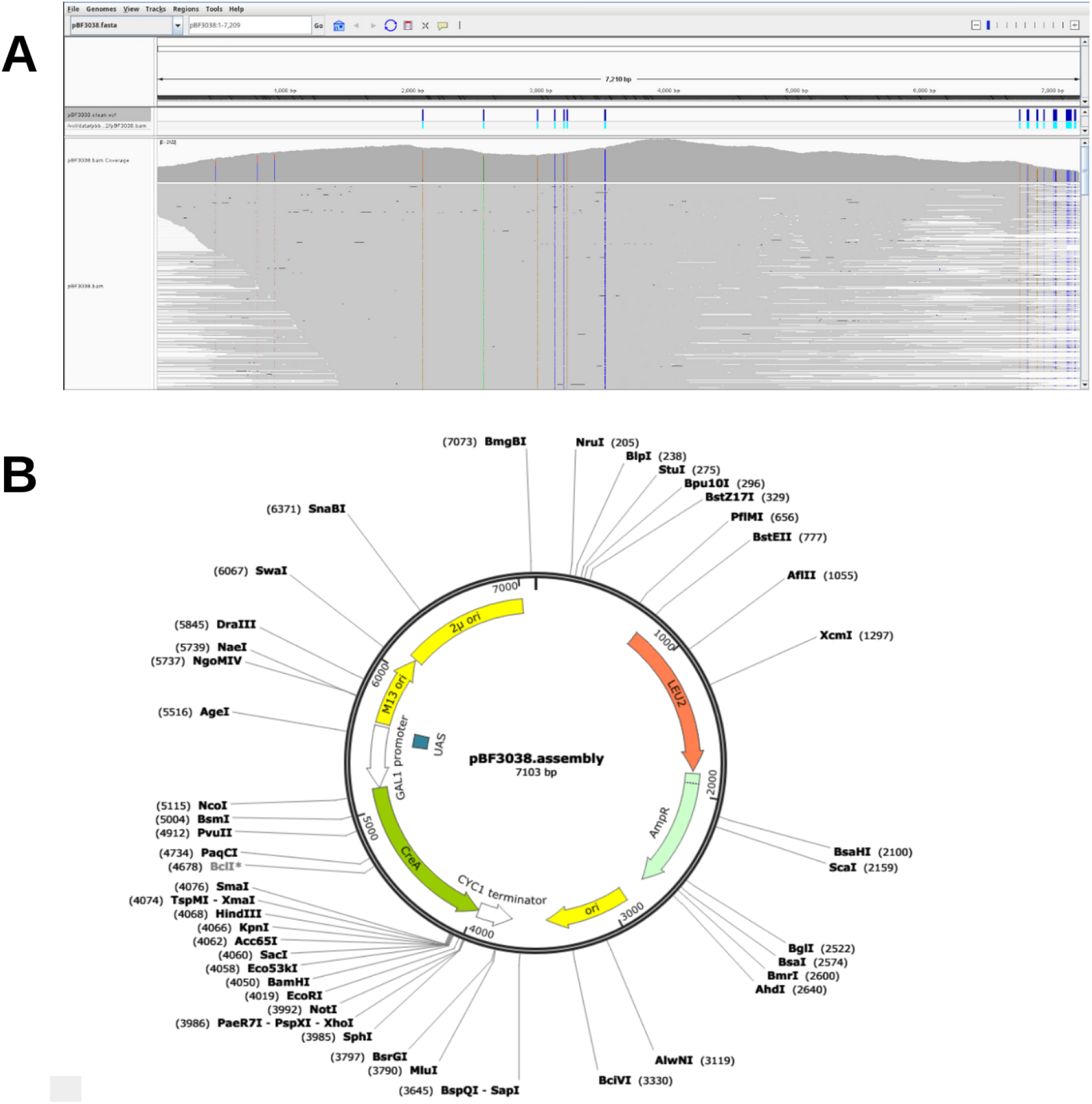
Screenshots of different results for pBF3038. A: Visualization of read mapping against the expected sequence of pBF3038 in IGV. Additionally, variant calling is included. B: Plasmid map of the pBF3038 sequence assembled with Miniasm.

The mapping of the assigned long reads shows that the entire length of the plasmid can be covered in single reads at a high coverage. Additionally, it also shows proof of concept that the reads can be assigned to the individual plasmids. Further, the resequencing and variant calling revealed several point mutations within our sequenced copy of pBF3038. An annotation of the assembled plasmid was able to validate the assembly process and showed that all key genetic elements of the plasmid were sequenced and assembled.

## Declarations

### Ethics approval and consent to participate

Not applicable

### Consent for publication

Not applicable

### Availability of data and materials

The scripts developed for this study are available via GitHub (https://github.com/bpucker/NanoPlasmiQC).

### Authors’ contributions

JAVSdO, VN, KW, and BP planned the project. VN and KW prepared the plasmid mixtures prior to library preparation. JAVSdO conducted library preparation, sequencing, and bioinformatic analyses. BP supervised the work, developed the tool for automatic data analysis, conducted bioinformatic analyses, and wrote the manuscript. All authors approved the final version of the manuscript and agreed to its submission.

## Acknowledgements

This work was supported by the de.NBI Cloud within the German Network for Bioinformatics Infrastructure (de.NBI) and ELIXIR-DE (Forschungszentrum Jülich and W-de.NBI-001, W-de.NBI-004, W-de.NBI-008, W-de.NBI-010, W-de.NBI-013, W-de.NBI-014, W-de.NBI-016, W-de.NBI-022). We thank all members of the Plant Biotechnology and Bioinformatics group for their support and feedback during the process.

